# 4-Methylumbelliferone Restores Age-Related Changes in Perineuronal Nets, Memory and Neuroinflammation

**DOI:** 10.64898/2026.01.30.702721

**Authors:** Anda Cimpean, Jana Dubisova, Noelia Martinez-Varea, Lucia Urdzikova-Machova, James W. Fawcett, Pavla Jendelova, Jessica C.F. Kwok

**Affiliations:** Department of Neuroregeneration, Institute of Experimental Medicine of the Czech Academy of Sciences, Prague, Czech Republic; Department of Clinical Neurosciences, John van Geest Centre for Brain Repair, University of Cambridge, Cambridge, United Kingdom; Faculty of Biological Sciences, University of Leeds, Leeds, United Kingdom

**Keywords:** aging, 4-methylumbelliferone, perineuronal nets, neuroinflammaging, neuroplasticity, cognitive decline

## Abstract

Aging leads to cognitive decline due to reduced neuronal plasticity and increased brain inflammation. Disruption of perineuronal nets (PNNs) is a recognized strategy to enhance neuroplasticity. Hyaluronan forms the backbone of PNNs, and 4-methylumbelliferone (4-MU), an inhibitor of hyaluronan synthesis, has been proposed to disrupt PNNs and enhance neuroplasticity in young rodents. In addition, 4-MU is also known as an immune response regulator, although its effects within the central nervous system (CNS) have been less characterized. In this study, we investigated the impact of long-term oral 4-MU administration in aged mice (20–22 months old), focusing on PNNs intensity, memory, and neuroinflammation. PNN intensity increased with age, and 4-MU treatment reduced it in 22-month-old mice to levels observed in 10-month-old animals; in the novel object recognition test, treated 22-month-old mice performed comparably to or better than 10-month-old controls. Moreover, markers of aging-associated neuroinflammation—including astrocytic and microglial activation as well as peripheral immune cell infiltration—were normalized to 10-month-old levels or further diminished following 4-MU treatment. Importantly, chronic 4-MU administration was well tolerated in aged mice, with no serious adverse effects observed. Together, these results suggest that 4-MU mitigates PNNs accumulation and neuroinflammaging while enhancing recognition memory, supporting its potential as a safe therapeutic approach for age-related cognitive decline.

## INTRODUCTION

Aging is characterized by a gradual decline in cognitive and motor functions, reducing the ability to learn, adapt to environmental changes, and recover from damage, impairments which are largely conserved across species. Aging is accompanied by a progressive loss of neuroplasticity within the central nervous system (CNS) (Brito et al., 2023). This reduction in plasticity reflects the convergence of multiple cellular, molecular, and structural processes. Among the key contributors are the accumulation of perineuronal nets (PNNs) (Karetko-Sysa et al., 2014; Foscarin et al., 2017; Mafi et al., 2020; Lehner et al., 2024) and an increase in neuroinflammation (Patterson, 2015; Tamatta et al., 2025).

PNNs are specialized extracellular matrix (ECM) structures that enwrap specific subgroups of neurons, most prominently parvalbumin-positive (PV+) inhibitory interneurons (Lupori et al., 2023). They form unique mesh-like pericellular coats that regulate the diffusion of neurotransmitters, ions, growth factors, and metabolites, as well as the distribution and clustering of synaptic receptors, thereby exerting a profound influence on synaptic transmission and neuronal function. PNNs are composed of a hyaluronan (HA) backbone anchored to the neuronal surface by hyaluronic acid synthase (HAS). Chondroitin sulfate proteoglycans (CSPGs) and tenascins attach to HA and are stabilized by link proteins (Crtl1/Hapln1, Bral2/Hapln4) (Testa et al., 2019; Auer et al., 2025). They are considered central pillars of neuroplasticity. Functionally, PNNs play a central role in closing critical periods of increased neuroplasticity in the CNS. Condensation of PNNs towards the end of neurodevelopment is highly linked to reduced synaptic flexibility, and impaired learning in aged subjects. Experimental degradation or remodeling of PNNs in adulthood has been shown to enhance neuroplasticity, resulting in increased associative, spatial, social, and auditory memory (Fawcett et al., 2022).

In parallel, brain aging is accompanied by profound changes in glial cells. These include altered morphology, increased numbers, and a shift toward reactive phenotypes. Within the tetrapartite synapse, glial reactivity and structural alterations disrupt synaptic homeostasis (Latham et al., 2023). Such changes contribute to reduced neuroplasticity, neuronal degradation, and increased vulnerability to neurodegenerative processes (Jurcau et al., 2024).

Aging is also associated with compromised blood–brain barrier (BBB) integrity, which facilitates T-cell infiltration into the CNS. Infiltrating immune cells interact bidirectionally with resident glia, creating a feedforward loop of inflammation: activated glia release pro-inflammatory cytokines (e.g., TNF-α, IL-1β, IL-6), chemokines (e.g., CCL2), reactive oxygen species (ROS, NO), and matrix metalloproteinases (MMP-2, MMP-9), which increase BBB permeability, further promoting T-cell entry and amplifying glial activation (Nevalainen et al., 2022; Zhang et al., 2022; Latham et al., 2023). Together, these changes contribute to “inflammaging”, a state of chronic low-grade inflammation that negatively affects synaptic function and neuroplasticity. Therapeutic strategies in this context aim to mitigate detrimental inflammatory cascades while preserving or enhancing protective glial functions.

The coumarin derivative 4-methylumbelliferone (4-MU) is a small molecule naturally occurring in plants of the Umbelliferae and Asteraceae families. Already approved in Europe and Asia for hepatoprotective, antispasmodic, and choleretic use (Nagy et al., 2015), 4-MU has gained attention for its capacity to modulate HA metabolism. Mechanistically, 4-MU depletes UDP-GlcA required for HA synthesis, downregulates hyaluronan synthase enzymes (HAS2) expression and upregulates hyaluronidase enzymes (Hyal1) expression, leading to an overall reduction in HA (Fedorova et al., 2025). Our previous studies have shown that 4-MU administration enhances neuroplasticity through PNN downregulation improving memory (Dubisova et al., 2022) and promotes serotonergic sprouting after spinal cord injury in rodents (Štepánková et al., 2023). Furthermore, 4-MU has well-established immunomodulatory properties, with studies demonstrating reduced inflammation in preclinical models of type 1 and type 2 diabetes, autoimmune arthritis, as well as lung, renal, and liver inflammatory pathologies (Fedorova et al., 2025). While its role in modulating immune responses within the CNS is less characterized, evidence indicates that 4-MU can attenuate glial scar formation after spinal cord injury (Štepánková et al., 2023), decrease immunoreactivity and disease progression in multiple sclerosis mice models (Kuipers et al., 2016b) and exert anti-inflammatory effects in rodent models of ischemic stroke (Mirshekari Jahangiri et al., 2024; Tamouk et al., 2025). These findings make 4-MU particularly relevant in the context of neuroinflammaging.

In this study, we set out to characterize age-related changes in PNNs and neuroinflammation, and to evaluate the effects of 4-MU on these processes. Our goal was to determine whether 4-MU treatment can enhance neuroplasticity by reducing PNNs and neuroinflammation, thereby enhancing recognition memory in aged mice. 4-MU reduced age-related PNN accumulation, normalized or further lowered neuroinflammation, and improved memory performance, highlighting its potential against cognitive decline in aging.

## MATERIALS AND METHODS

### 1. Animals

We used three groups of C57BL/6JOlaHsd mice. The “10m” group (n = 5) entered the study at 4 months of age and was fed chocolate-flavored chow (Sniff GmbH, Soest, Germany) (placebo) for 6 months, reaching 10 months of age at the end of the experiment. The “22m” group entered the study at 14–16 months of age and was divided into two subgroups: control (n = 15) and treated (n = 15). The control subgroup received chocolate-flavored chow, while the treated subgroup received chocolate-flavored chow supplemented with 5% (w/w) 4-MU (4-methylumbelliferone, DbPharma France) corresponding to a daily dose of 36.9 µmol/g for 6 months. At the end of the experiment, the old group was 20–22 months old. All procedures were conducted in compliance with the European Communities Council Directive of 22 September 2010 (2010/63/EU), adhered to the ARRIVE guidelines, and were approved by the Ethics Committee of the Institute of Experimental Medicine CAS, Prague, Czechia. Animals had ad libitum access to food and water and were maintained on a 12 h light/dark cycle (lights off at 7:30 p.m.). Behavioral testing was performed during the light phase. At the end of the treatment period, animals underwent behavioral assessment of memory and motor functions, were weighed, and then sacrificed for blood and urine collection and brain isolation. Blood was collected by cardiac puncture into BD Microtainer® tubes. Samples for hematological analysis were drawn into K_2_EDTA-coated tubes (#365974), and the following parameters were measured: leukocytes, erythrocytes, platelets, reticulocytes, neutrophils, lymphocytes, monocytes, eosinophils, and basophils. For biochemical analyses, blood was collected in serum-separating tubes (#365963) and urine samples were obtained via bladder puncture to determine glucose and bile acid levels. All measurements were performed by Synlab (Munich, Germany).

### 2. Behavioral tests

To evaluate the effects of 4-MU on aging, a series of behavioral tests were performed. Memory performance was assessed using the spontaneous object recognition (SOR) task, while motor function was evaluated using the rotarod and grip strength tests, which probe coordination, joint mobility, and muscle integrity.

#### 2.1 Spontaneous object recognition task (SOR)

The SOR task was performed in a Y-maze, in which one arm served as the start arm (4.5 cm long, 8 cm wide) and the other two arms (10 cm long, 8 cm wide) were used for object presentation. Randomly shaped junk objects (∼10 × 5 × 5 cm) were used as stimuli. All mice were habituated to the maze in two daily sessions prior to testing. During the sample phase, mice were allowed to explore two identical objects for 5 min. After a delay of 6 or 24 h in the home cage, the choice phase was conducted. This phase was procedurally identical to the sample phase, except that one of the familiar objects was replaced with a novel object. Mice were again allowed to explore for 5 min. Behavioral performance was video-recorded and analyzed offline by an experimenter blinded to treatment. Exploration time was defined as the duration during which the mouse oriented toward and contacted the object. Instances in which animals climbed or sat on objects were excluded. A discrimination score was calculated as (time exploring the novel object – time exploring the familiar object) / (total exploration time). Thus, a score of 1 indicated exclusive exploration of the novel object, while a score of 0 indicated equal exploration of both objects. Statistical analysis was performed using one-way ANOVA followed by Tukey’s multiple comparisons test.

#### 2.2 Rotarod and grip test

Motor coordination and balance were assessed in the old group using a rotarod apparatus equipped with automatic timers and fall sensors (ROTA-ROD 47700, UGO BASILE S.R.L., Italy) with a 7 cm diameter drum. Prior to testing, mice were habituated to remain on the stationary drum for 60 s and then pre-trained at a constant speed of 5 rotations per minute (RPM) for 120 s. During the test phase, animals were tested at 10 RPM for up to 300 s in three consecutive daily sessions following training. The latency to fall was automatically recorded. Muscle strength was assessed using a grip strength meter (BIO-GS3, Bioseb, Vitrolles, France). Mice were tested for hind limb grip strength in three consecutive daily sessions. Each mouse was allowed to grasp the metal grid with its hind paws and was gently pulled backward until the grip was released; the maximal force exerted was recorded automatically. Statistical analysis was performed using unpaired t-test.

### 3. Tissue preparation and immunohistochemistry

Mice were deeply anesthetized with an intraperitoneal injection of an overdose of pentobarbital (Sigma-Aldrich, St. Louis, MO, USA) and intracardially perfused with phosphate-buffered saline (PBS), followed by 4% paraformaldehyde (PFA) in 0.1 M phosphate buffer. Heads were post-fixed in 4% PFA for 1 week. Brains were then dissected from the skull and cryoprotected in increasing concentrations of sucrose (10–30%, w/v). After cryoprotection, brains were embedded in O.C.T. compound (00411243, VWR), frozen, and coronally sectioned at 20 μm thickness using a cryostat (CryoStar NX70, Thermo Fisher Scientific, MA, USA). Sections were collected free-floating in PBS and stored at –20 °C in cryoprotectant solution until use. For immunohistochemistry, free-floating sections were permeabilized with 0.5% (v/v) Triton X-100 in PBS for 20 min at room temperature. Lipofuscin autofluorescence was quenched using 50 mM ammonium acetate buffer with CuSO_4_. Endogenous biotin was blocked using an endogenous biotin blocking kit (ab64212, Abcam, Cambridge, UK). Perineuronal nets (PNNs) were visualized using Wisteria floribunda agglutinin (WFA), which binds N-acetylgalactosamine β1 residues of glycoproteins within the neuronal extracellular matrix (1:150; biotinylated WFA, L1516, Sigma-Aldrich). Non-specific binding of WFA was blocked with 10% Chemiblocker (#2170, Merck, Darmstadt, Germany) in PBS containing 0.2% Triton X-100 for 2 h at room temperature. Sections were incubated overnight at 4 °C with primary antibodies against GFAP (chicken, 1:800, ab4674, Abcam), Iba1 (goat, 1:800, 019-19741, Fujifilm), CD45 (rabbit, 1:300, bs-10599R, Bioss), and biotinylated WFA. Nuclei were counterstained with DAPI (1:1000, D1306, Invitrogen). The following secondary antibodies were used: streptavidin–Alexa Fluor 488 (1:200, S32354, Life Technologies), donkey anti-rabbit Alexa Fluor 594 (1:400, A-31556, Thermo Fisher Scientific), goat anti-chicken Alexa Fluor 488 (1:400, A-11039, Thermo Fisher Scientific), and donkey anti-goat Alexa Fluor 488 (1:400, A11055, Invitrogen). After staining, sections were mounted on glass slides, coverslipped with antifade mounting medium, and stored at 4 °C until imaging.

### 4. Imaging and analysis

Images were acquired using an Olympus SpinSR10 inverted fluorescence confocal spinning disk microscope. A 20× objective was used for general imaging, and a 40× objective was used for glia-morphology analysis. WFA-labeled perineuronal nets (PNNs) were imaged in the cerebral cortex and hippocampus, while inflammation-related analyses were focused on the somatosensory cortex, corpus callosum and hippocampus. Image analysis was performed using ImageJ (NIH) software. For PNN quantification, fluorescence signal was measured using the Mean Intensity of Top Quartile Pixels method. Briefly, for each image, the top 25% of brightest pixels within the region of interest were identified, and their average fluorescence intensity was calculated (arbitrary units, a.u.). This approach reduces variability caused by background staining and highlights the most strongly labeled PNN structures. Astrocytes were analyzed by GFAP immunostaining. The total number of GFAP^+^ cells and the mean GFAP signal intensity were quantified. Astrocytic morphology in the somatosensory cortex and hippocampus was assessed using Sholl analysis, in which concentric circles of increasing radius are drawn around the cell soma and the number of intersections between processes and circles is counted (starting radius: 3 µm; step size: 3 µm). This method provides a measure of astrocytic branching complexity and arborization. In the corpus callosum, this analysis could not be performed due to the elongated morphology of astrocytes in this region. Instead, we quantified the cell body area, as soma hypertrophy is a well-established feature of reactive astrocytes. Microglia were examined by Iba1 immunostaining. Similar to astrocytes, we quantified the total number of Iba1^+^ cells, the mean Iba1 fluorescence intensity, and microglial morphology using Sholl analysis (starting radius: 5 µm; step size: 3 µm). To evaluate peripheral immune cell infiltration, a combination of Iba1 and CD45 staining was used. Since CD45 labels both microglia and peripheral immune cells, cells positive for CD45 but negative for Iba1 were manually counted by a blinded experimenter. Statistical analyses were performed using ordinary one-way ANOVA followed by Tukey’s multiple comparisons test.

## RESULTS

### 1. 4-MU treatment reverses age-related increase in PNNs and enhances object recognition memory in aged mice

In our previous work, we showed that oral 4-MU treatment in 10-months old adult mice (10m) reduced total CNS glycosaminoglycan levels, decreased PNN formation, modulated the expression of HA- and PNN-related genes, and improved object recognition memory (Dubisova et al., 2022). In the present study, we sought to assess how PNNs change with aging and to evaluate the impact of long-term 4-MU treatment on these age-related alterations.

We performed WFA staining for PNNs on coronal brain sections from 10 months old adult (10m) mice, 22-months-old aged mice (22m), and 22m aged mice treated with 5% (w/w chow) 4-MU for 6 months (22m+4MU). Our analysis focused on the somatosensory cortex and hippocampus, regions critical for cognitive function (Fig. 1A). The results revealed that aging significantly increased WFA intensity in both cortex and hippocampus when compared to 10m adult (22m cortex: 0.42 ± 0.44 vs. 10m cortex: 0.04 ± 0.035, p=0.0337; 22m hippocampus: 0.53 ± 0.26 vs. 10m hippocampus: 0.14 ± 0.04, p=0.0058). 4-MU treatment however showed comparable WFA intensities to 10m adult mice and significantly lower than their age-matched controls (22m+4MU cortex: 0.06 ± 0.08, p=0.0105; 22m+4MU hippocampus: 0.2 ± 0.085, p=0.0177), demonstrating that 4-MU treatment effectively reduced PNN intensity and restored it to young adult levels (Fig. 1B,C).

**Fig. 1:**
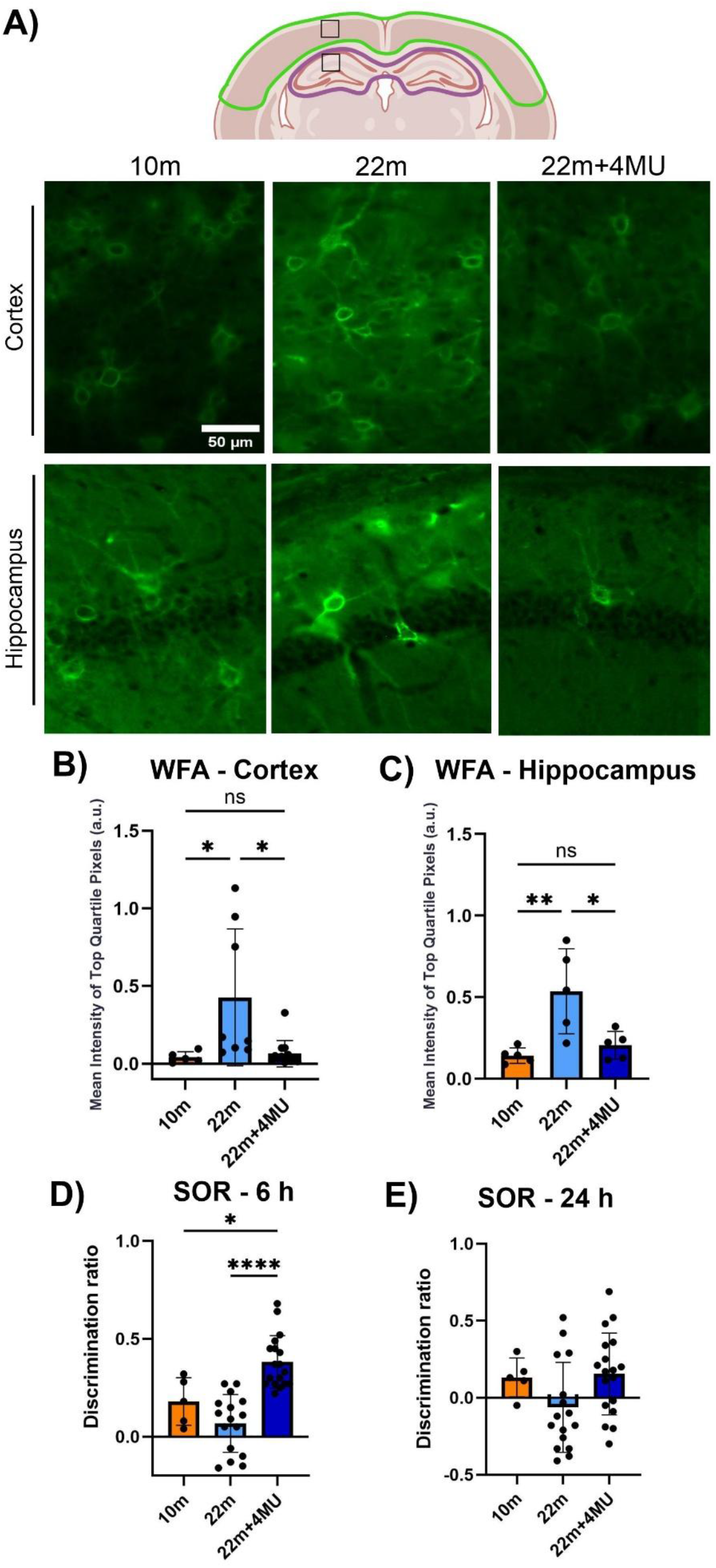
4-MU reverses age-related PNN accumulation and improves memory performance. (A) Schematic brain representation of the areas studied and representative images of WFA immunofluorescence in cerebral cortex and hippocampus of 10-months-old (10m), 22-months-old (22m) and 22-months-old mice treated with 4-MU for 6 months (22m+4MU). (B,C) Quantification of WFA intensity in cortex and hippocampus, respectively, showing increase of PNN intensity with aging and reduction after 4-MU treatment. (D,E) SOR performance at 6 h and 24 h retention intervals, respectively. Data are presented as mean ± SD and were analyzed using ordinary one-way ANOVA; *p < 0.05, **p < 0.01, **** p < 0.0001.

To examine whether this reduction influenced memory performance in aged mice, we evaluated spontaneous object recognition (SOR) memory in the three groups of mice. Memory was tested at 6 h and 24 h delays between familiarization and testing, intervals that require consolidation, with 24 h representing a benchmark for robust long-term memory in rodents (Villar et al., 2017; Cinalli Jr. et al., 2020). As expected, aged mice performed worse than young adults at both intervals, though the differences did not reach significance (discrimination ratios average: 10m 6 h = 0.18 ± 0.12 vs. 22m 6 h = 0.068 ± 0.14, p=0.27; 10m 24 h = 0.13 ± 0.12 vs. 22m 24 h = –0.06 ± 0.29, p=0.33). 4-MU–treated aged mice showed improved performance at both delays. At 6 h, treated aged mice outperformed both young and old groups (discrimination index average 22m+4MU: 0.38 ± 0.13, p=0.0178 compared to 10m; p<0.0001 compared to 22m) (Fig 1D). At 24 h, while not statistically significant, a clear trend toward improvement was observed (discrimination index average 22m+4MU: 0.15 ± 0.26, p=0.33 compared to 10m; p=0.057 compared to 22m) (Fig 1E). Together, these findings indicate that oral 4-MU treatment reverses age-associated PNN accumulation and alleviate aspects of age-related cognitive decline.

### 2. Ageing-induced astrocytic activation is suppressed by 4-MU treatment

We next investigated the effect of 4MU on chronic inflammation in aging. Given that neuroinflammaging has been reported to predominantly affect white matter regions (Raj et al., 2017; Hahn et al., 2023), we also included the corpus callosum in our assessment of neuroinflammatory changes in addition to the somatosensory cortex and hippocampus (Fig. 2A). Our analysis revealed region-specific patterns of astrocytic changes with aging (Fig. 2B). In control 22m aged animals, there is no difference in the number of GFAP positive cells in the somatosensory cortex and hippocampus, but we observe a significant increase in the corpus callosum when compared to 10m controls, whereas aged animals treated with 4-MU displayed values comparable to young controls (Fig. 2C). Similarly, GFAP intensity tended to increase in the somatosensory cortex and was significantly elevated in the corpus callosum and hippocampus. Notably, 4-MU–treated 22m aged animals showed reduced GFAP intensity across all regions compared with both aged and young controls (Fig. 2D).

**Fig. 2:**
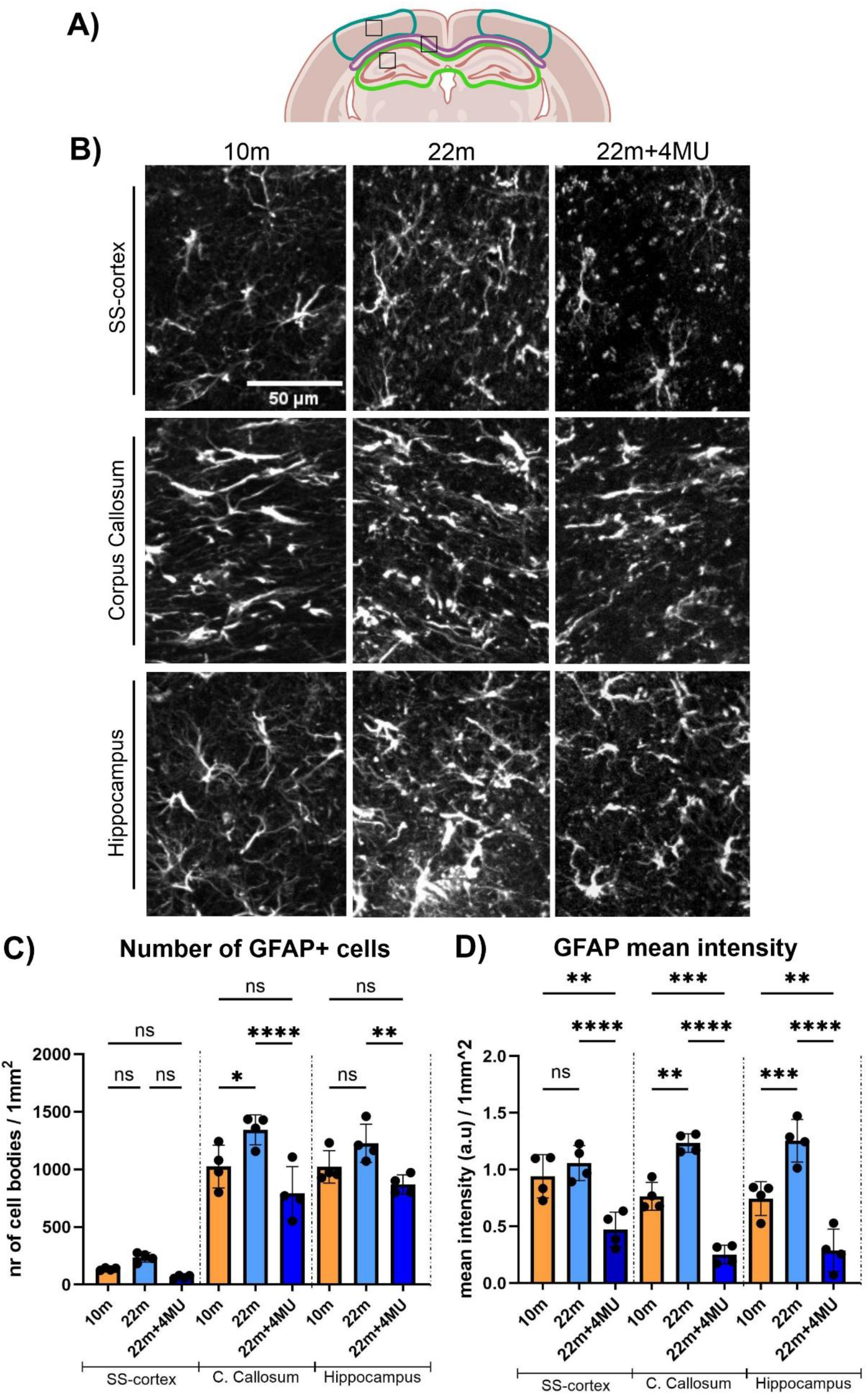
Astrocytes cell number and mean fluorescence intensity in cortex, corpus callosum, and hippocampus. (A) Schematic representation of the brain regions analyzed for neuroinflammatory changes. (B) Representative GFAP immunofluorescence images in the indicated areas. (C,D) Quantification of GFAP^+^ cell density and mean fluorescence intensity, respectively, showing increased astrocytic activation with age reversed by 4-MU. Values are mean ± SD; ordinary one-way ANOVA; *p < 0.05, **p < 0.01, *** p < 0.001, **** p < 0.0001.

Morphological analysis further demonstrated a significant reduction in astrocytic arborization in aged animals within the somatosensory cortex and hippocampus (Fig. 3A–C), along with soma hypertrophy in the corpus callosum (Fig. 3D,E). The alterations in the somatosensory cortex and hippocampus were largely normalized by 4-MU treatment. Together, these findings indicate that aging induces region-specific astrogliosis, with white matter regions being particularly vulnerable, showing increased number of GFAP positive cells, GFAP intensity and soma hypertrophy. Treatment with 4-MU effectively counteracted these changes, revealing a strong astrocytic immunomodulatory effect and suggesting its potential to mitigate aging-associated astrogliosis.

**Fig. 3:**
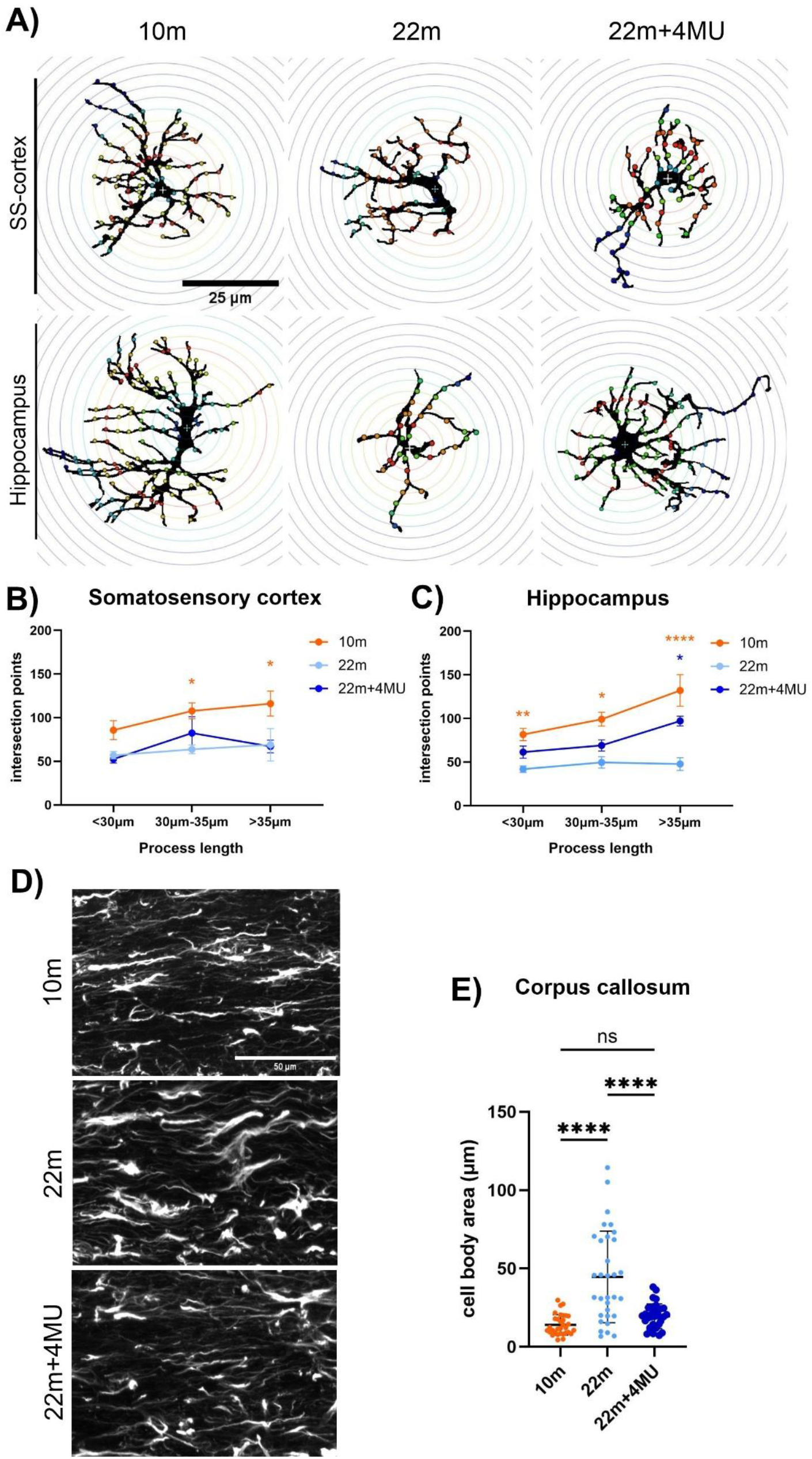
Morphological analysis of astrocytes. (A) Representative images of individual GFAP+ astrocytes assessed using the Scholl method in the somatosensory cortex and hippocampus. (B,C) Quantification of the number of intersection points per cell, demonstrating a reduction in ramification with age, with 4-MU-treated older animals showing values similar to young controls. (D) Representative images of astrocytes in the corpus callosum, and (E) quantification of soma area across the three animal groups, illustrating an increase in astrocyte cell body size with aging, while 4-MU treatment significantly reduces this area, bringing values closer to those seen in young animals. Data are presented as mean ± SD; analyzed by one-way ANOVA; *p < 0.05, **p < 0.01, **** p < 0.0001.

### 3. Aging-induced microglia activation is restored by 4-MU treatment

Another major contributor to neuroinflammaging is microglial reactivity (Von Bernhardi and Eugenín, 2025). We next examined whether 4-MU treatment could counteract aging-associated microglial alterations. We therefore analyzed Iba1 positive cell density, Iba1 signal intensity (Fig. 4A), and performed a morphological assessment of microglial arborization (Fig. 5A) in the somatosensory cortex, hippocampus and corpus callosum. In contrast to the astrocytic changes described above, aging induced a robust and significant increase in both Iba1 positive cell number (Fig. 4B) and Iba1 intensity (Fig. 4C) across all brain regions studied. 4-MU treatment significantly decreased the density of Iba1 positive cells and Iba1 intensity to levels comparable to 10m adults. Moreover, microglial morphology was also altered, as revealed by Sholl analysis: 22m aged mice exhibited significantly reduced arborization compared to 10m adults, consistent with a transition toward a reactive phenotype in the brain regions studied. 4-MU treated old animals in contrast showed microglial arborization comparable to 10m controls (Fig. 5 B-D).

**Fig. 4:**
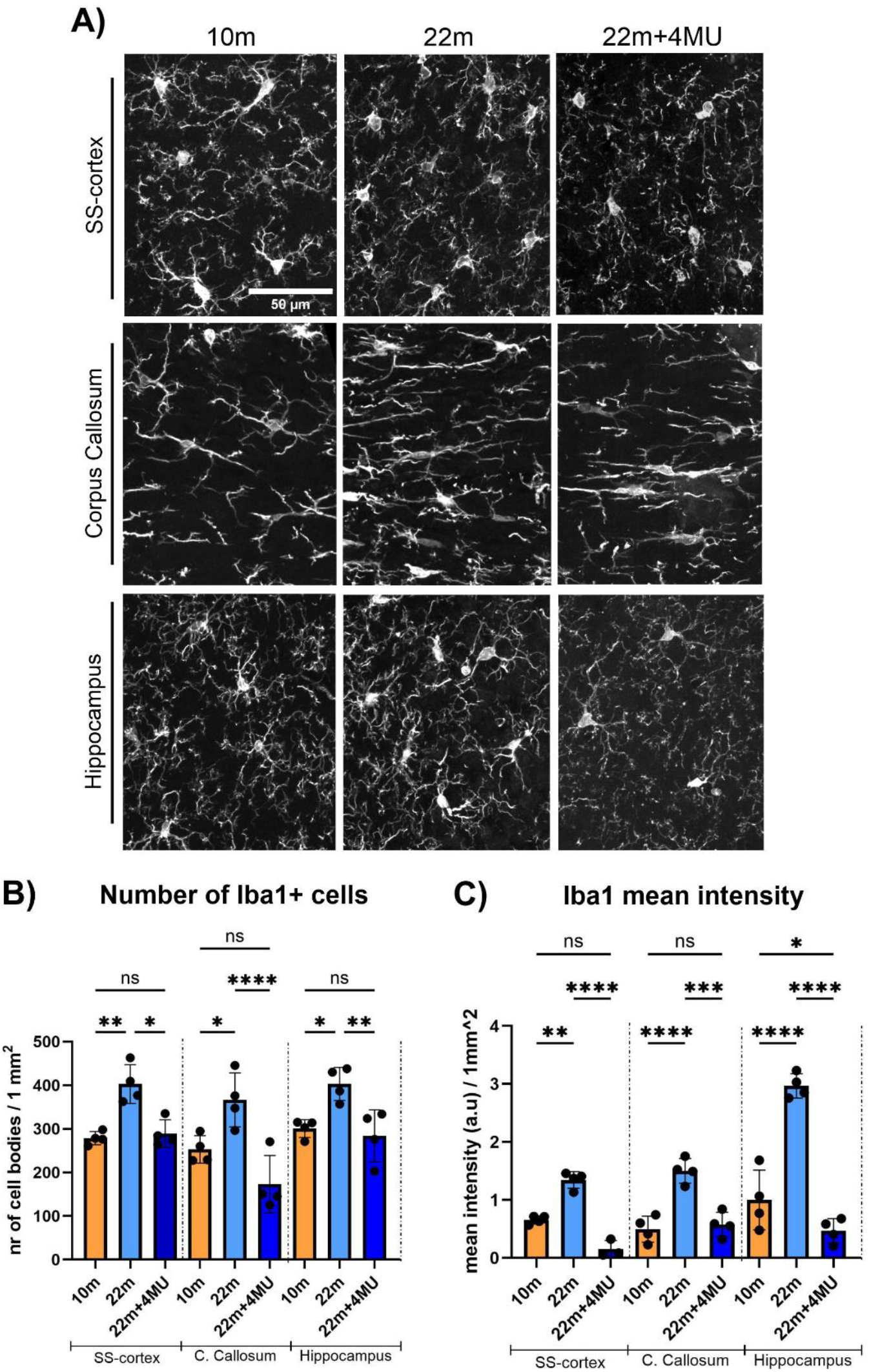
Microglia cell number and mean fluorescence intensity across brain regions. (A) Representative Iba1 staining in the somatosensory-cortex, corpus callosum and hippocampus. (B,C) Quantification of Iba1^+^ cell density and mean fluorescence intensity, respectively, revealing microglial activation in aged mice attenuated by 4-MU. Data are mean ± SD; ordinary one-way ANOVA; *p < 0.05, **p < 0.01, *** p < 0.001, **** p < 0.0001.

**Fig. 5:**
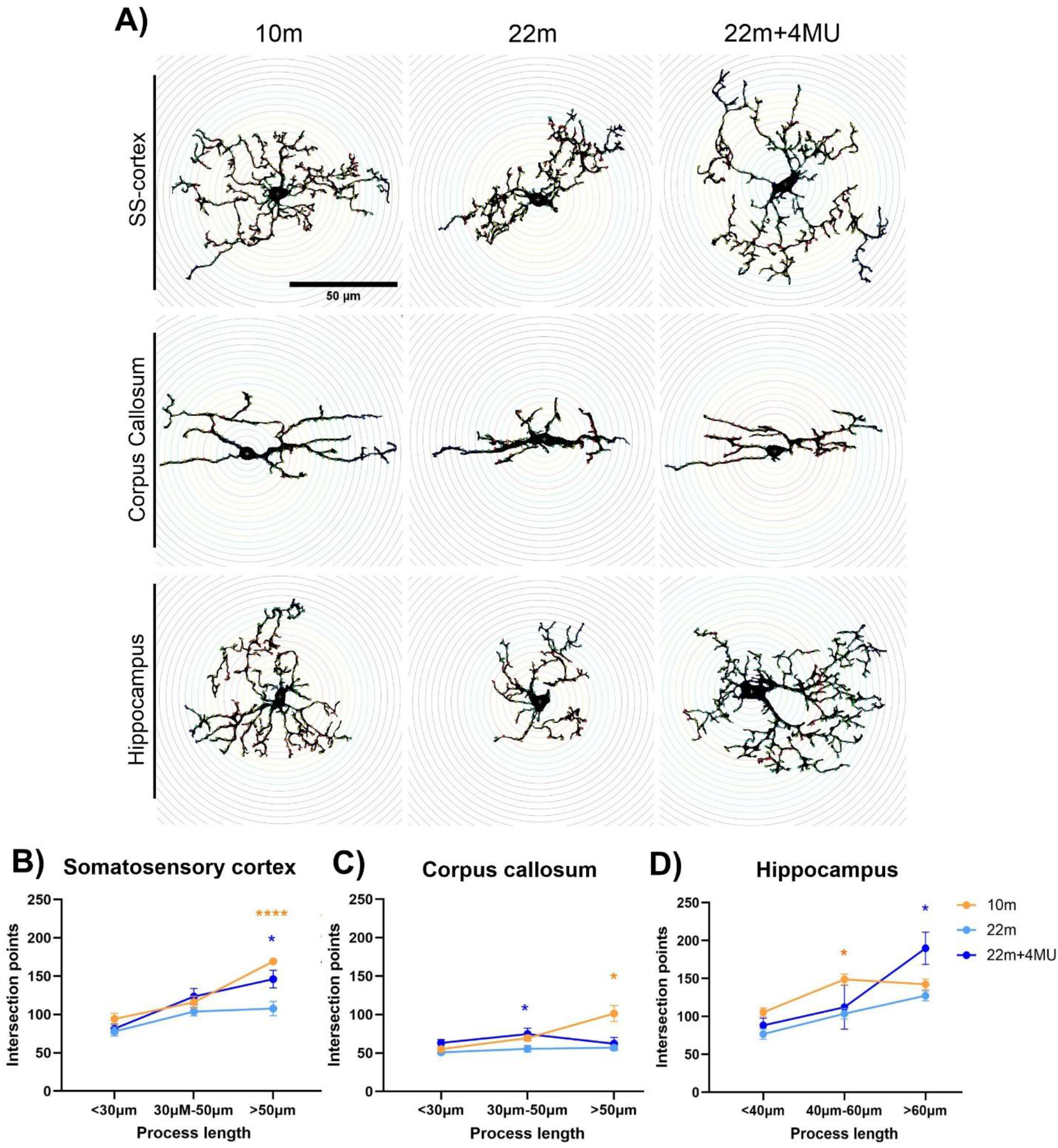
Morphological analysis of microglia. (A) Representative images of individual IBA1^+^ microglia used for morphological assessment. (B-D) Quantification of the number of intersection points per cell in the somatosensory-cortex, corpus callosum and hippocampus, indicating reduced ramification with age and restoration by 4-MU. Values are presented as mean ± SD; ordinary one-way ANOVA; *p < 0.05, **** p < 0.0001.

Together, these findings suggest that 4-MU alleviates key age-related microglial alterations across all examined regions, consistently reducing activation and restoring arborization, indicative of a broad modulatory influence on microglial remodeling with age.

### 4. Peripheral immune cell infiltration increases in aged brains and is normalized by 4-MU

With aging, the BBB becomes more permeable, facilitating infiltration of peripheral immune cells. HA has been implicated in immune cell recruitment, and 4-MU has been reported to reduce immune cell infiltration across multiple organs (McKallip et al., 2015; Suarez-Fueyo et al., 2019; Bosi et al., 2022; Galkina et al., 2022). In the CNS, this phenomenon is less well characterized. Existing evidence suggests that 4-MU can also limit infiltration, but only in disease or injury models (Mueller et al., 2014; Kuipers et al., 2016b, pp. 4-; Štepánková et al., 2023), not under normal aging conditions where BBB alterations follow different dynamics. To address this gap, we examined peripheral immune cell infiltration into the brains of 10m adult, aged, and aged 4-MU–treated animals. We quantified infiltrating immune cells using CD45 as a pan-leukocyte marker. Because CD45 is also expressed by microglia, we combined CD45 immunolabeling with Iba1 to identify peripheral immune cells as CD45-positive/Iba1–negative. Analyses were performed in the somatosensory cortex, corpus callosum, and hippocampus (Fig. 6A). Results showed a significant increase in CD45-positive/Iba1-negative cells in 22m aged mice compared with 10m controls in the corpus callosum, while the somatosensory cortex and hippocampus showed no substantial changes. These findings highlight that age-related immune infiltration is most prominent in white matter regions, consistent with other hallmarks of neuroinflammaging. Importantly, 4-MU treatment markedly reduced the elevated immune cell numbers in the corpus callosum, restoring them to levels observed in 10m animals (Fig. 6B). Our results indicate that 4-MU can reduce age-related peripheral immune cell infiltration into the corpus collosum, which may limit inflammatory signaling, thereby creating a more permissive environment for neuronal health and neuroplasticity.

**Fig. 6:**
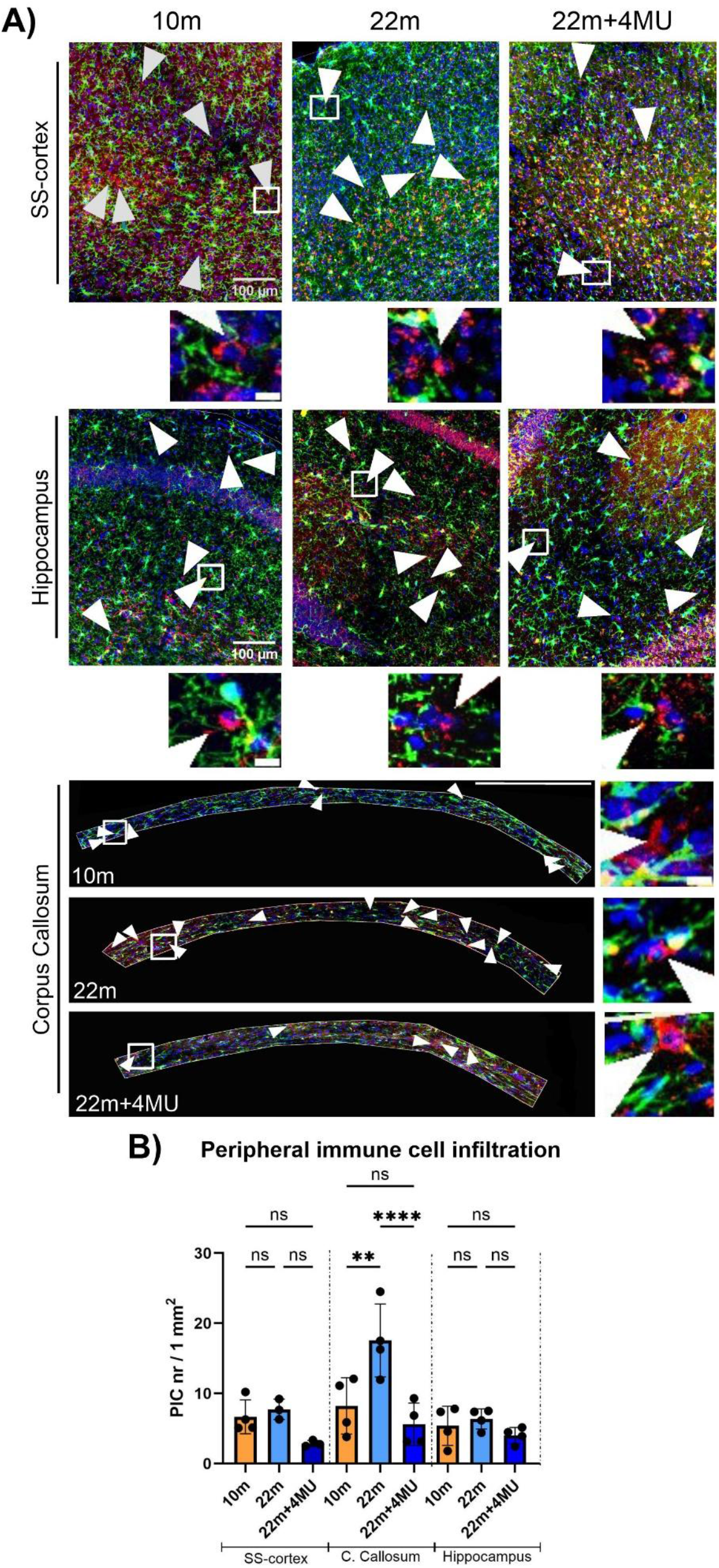
Peripheral immune cell (PIC) infiltration in cortex, corpus callosum and hippocampus. (A) Representative images of CD45 (red), Iba1 (green) and DAPI (blue) staining in the indicated regions. Arrows mark CD45^+^Iba1^−^ cells and the insets show a magnified view of one such cell. (B) Quantification of CD45^+^Iba1^−^ cells per mm^2^ in the indicated areas, showing increased number of PICs in the corpus callosum with aging, which was reduced to young adult levels after 4-MU treatment. Data are presented as mean ± SD and were analyzed using ordinary one-way ANOVA; **p < 0.01, **** p < 0.0001.

### 5. Long-term oral 4-MU treatment is well tolerated in aged mice

Many previous studies have described 4-MU as a safe compound with minimal side effects, although most available data derive from 3–6 months of administration (Yates et al., 2015; Dubisova et al., 2022; Štěpánková et al., 2023). Notably, one study reported lifelong treatment with 4-MU in mice without observing major adverse effects (Nagy et al., 2024). In addition, its wash-out effect has been characterized, showing that any minor side effects do not persist after treatment cessation (Dubisova et al., 2022; Štěpánková et al., 2023). However, given that our study involved old mice, whose physiology may differ from animals in previous studies, we evaluated multiple parameters to assess 4-MU safety in aged animals (Fig. 7). The body weight of the animals showed a significant reduction in 4-MU– treated mice compared to controls, similar to previous studies (Fig. 7A). As HA is a key component for cartilage functions, motor and joint mobility were assessed by grip strength (Fig. 7B) and rotarod tests (Fig. 7C), and found no differences between treated and 22m control mice. Hematological analyses, including total leukocytes, erythrocytes, platelets, reticulocytes, and leukocyte subpopulations (neutrophils, lymphocytes, monocytes, eosinophils, and basophils), showed values within the expected physiological range, with no differences between treated and control mice (Fig. 7D). Finally, we assessed bile acids and glucose levels in blood and urine, parameters previously reported to be influenced by 4-MU treatment. Although no significant changes were detected, a trend toward increased levels exceeding the physiological range (Unno et al., 2023; Hönes, 2024; Norman et al., 2024; Zheng et al., 2024) was observed, particularly in glycosuria (Fig. 7E–H). Taken together, these results indicate that long-term oral administration of 4-MU is generally safe in old mice, causing no major hematological, biochemical, or functional alterations.

**Fig. 7:**
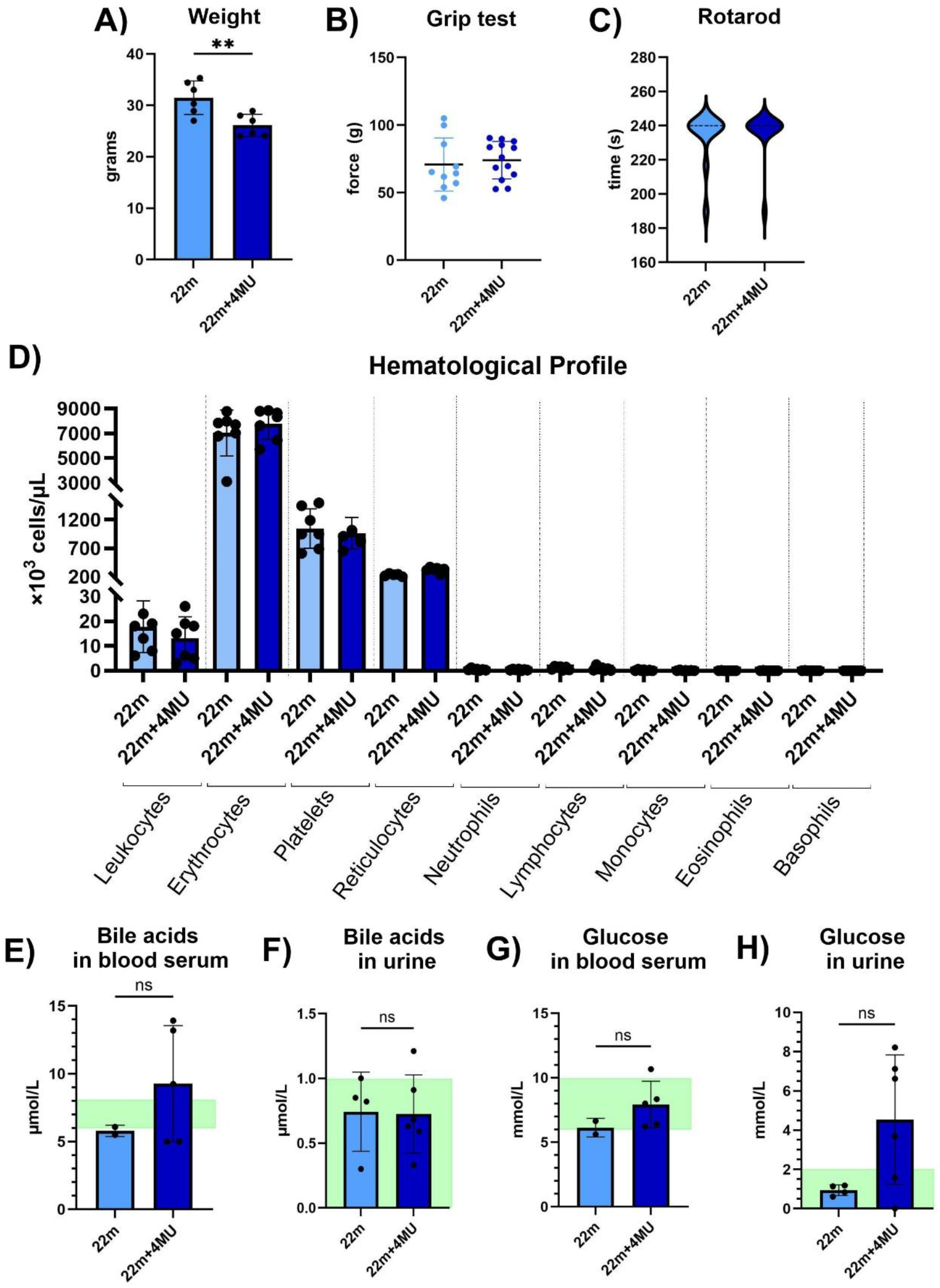
Assessment of safety parameters in aged mice following long-term 4-MU treatment. (A) Body weight measured at the end of the experimental period. (B) Forelimb grip strength. (C) Rotarod performance. (D) Hematological profile. (E-H) Bile acids and glucose concentrations in blood and urine, respectively. Physiological range is indicated in green. Data are presented as mean ± SD. Statistical comparisons between old control and old 4-MU–treated groups were performed using unpaired t-test with significance indicated as ** p < 0.01.

## DISCUSSION

Here, we show that aging robustly increases PNN intensity in the regions examined. In addition, it also led to region-specific neuroinflammatory changes in astrocytes, microglia, and infiltrating peripheral immune cells. Chronic 6 months 4-MU treatment reversed these alterations and improved object recognition memory, suggesting complementary effects of PNN modulation and reduced neuroinflammation.

The ECM plays key regulatory roles in neuronal and glial function through diverse and dynamic interactions. HA, a major glycosaminoglycan distributed across grey and white matter, signals through receptors such as CD44, RHAMM (HMMR), LYVE-1, ICAM-1, and TLR2/4 to influence proliferation, differentiation, migration, activation states, and process extension in stem cells, neurons, astrocytes, and microglia (Peters and Sherman, 2020). HA also forms the structural backbone of PNNs, specialized ECM assemblies that support circuit stability and excitatory–inhibitory balance. Furthermore, HA contributes to endothelial barrier regulation and lymphocyte extravasation (Galkina et al., 2022), and accordingly, modulation of HA metabolism has gained increasing interest in CNS disorders. Previous studies have reported age-related increases in PNN density in rodents (Karetko-Sysa et al., 2014; Mafi et al., 2020) and in humans (Lehner et al., 2024), accompanied by changes in sulfation patterns that render PNNs more inhibitory (Foscarin et al., 2017). Consistent with this literature, we observed a marked increase in PNN intensity in both cortex and hippocampus of aged animals. Chronic 4-MU treatment restored PNN intensity to levels comparable to young controls, suggesting that 4-MU can counteract age-associated PNN remodeling. PNNs are central regulators of neuroplasticity and have been implicated in the closure of critical periods, ocular dominance plasticity, auditory relearning, fear extinction, and multiple forms of memory (Reichelt et al., 2019). In our previous work, we showed that oral 4-MU reduces PNN intensity and enhances object recognition memory in young adult mice (Dubisova et al., 2022). Experimental PNN degradation using chondroitinase-ABC, which digests CSPGs, substantially prolongs long-term object recognition memory in both wild-type and MAPT mouse models of dementia (Romberg et al., 2013; Yang et al., 2015). Similarly, mice lacking Crtl1/Hapln1 in the CNS display markedly extended recognition memory compared with controls (Romberg et al., 2013). Age-related cognitive decline primarily affects the acquisition of new long-term memories, while retrieval of remote memories remains largely preserved (Brito et al., 2023). Because the SOR task specifically evaluates acquisition of new information, it is well suited for aging studies. Notably, aged mice treated with 4-MU showed enhanced object recognition, surpassing young controls at the 6-hour interval, and approaching young performance at 24 hours. These results are consistent with previous literature and suggest that modulating HA metabolism may represent a promising strategy to mitigate age-related cognitive impairment.

Regarding neuroinflammaging, transcriptomic analyses of the aged human brain have shown that glial cells, rather than neurons, undergo the majority of age-associated transcriptional changes (Soreq et al., 2017). Astrocytes participate in the tetrapartite synapse, support neurons, interact with vasculature, and contribute to BBB integrity. Microglia serve as the innate immune cells of the CNS and, beyond their injury response, have essential homeostatic roles including regulation of synapse formation, maturation, and elimination, as well as modulation of neurogenesis, myelination, and neuroplasticity. With aging, both cell types show reduced homeostatic capacity and elevated inflammatory reactivity, increasing neuronal vulnerability and predisposing the brain to age-related decline. Region-specific glial changes have been described, typically characterized by stable or increased cell numbers, elevated marker expression, hypertrophic somata, and reduced ramification in both astrocytes (Clarke et al., 2018; Cotto et al., 2019) and microglia (Raj et al., 2017; Ana, 2024), though, with distinct patterns across brain regions (Salas et al., 2020). In line with the literature, we observed the following aging-related alterations:

- Somatosensory cortex: Minimal neuroinflammatory changes. Astrocyte numbers and GFAP expression remained stable, although arborization was reduced. Microglia displayed increased numbers, elevated Iba1 expression, and reduced branching.
- Hippocampus: Moderate neuroinflammatory changes. Astrocyte numbers were unchanged, but GFAP expression increased and arborization decreased. Microglia exhibited elevated numbers and marker expression with reduced arborization.
- Corpus callosum: The most pronounced alterations. Astrocyte number and GFAP expression were significantly increased, with hypertrophic morphologies consistent with reactivity. Microglia showed robust increases in number and activation markers and substantial reductions in arborization.

These results mirror previous observations that white matter regions are particularly vulnerable to neuroinflammaging (Raj et al., 2017; Hahn et al., 2023), that the hippocampus exhibits more pronounced changes than neocortical areas such as the somatosensory cortex (Salas et al., 2020), and that microglia display more marked aging-associated alterations than astrocytes, including in non-human primates (Salas et al., 2020). Chronic 4-MU treatment in aged mice effectively reversed these neuroinflammatory signatures across all examined regions, restoring astrocytic and microglial parameters to levels comparable to young controls. HA regulates glial activation via TLR4, and several *in vitro* (Chistyakov et al., 2020) and *in vivo* (Kuipers et al., 2016b; Štepánková et al., 2023) studies have shown that 4-MU attenuates HA-dependent inflammatory responses, supporting our observations. Aging is also associated with BBB deterioration and increased permeability. Tissue-resident T cells have been described to accumulate in the white matter of older individuals (Jurcau et al., 2024). In our study, the corpus callosum was the only region displaying infiltration of peripheral immune cells. HA contributes to immune cell recruitment by binding CD44 on vascular endothelium facilitating leukocyte adhesion, and HA fragments signal through TLR2/4 to promote inflammatory gene expression. 4-MU has been shown to reduce immune cell infiltration in multiple organs (McKallip et al., 2015; Suarez-Fueyo et al., 2019; Bosi et al., 2022) and in the CNS during EAE, consistent with our finding that 4-MU reduces peripheral immune cells infiltration in the aging brain. Together, these results suggest that 4-MU exerts broad immunomodulatory effects and mitigates age-related neuroinflammation, potentially acting in concert with reduced PNN density to support cognitive improvement. 4-MU may also influence other cell types, such as stem cells, and additional brain regions that we have not examined, which underscores the need for further research.

Regarding safety, previous studies reported no major adverse effects of 4-MU and reversibility of observed changes after washout (Kuipers et al., 2016a; Štěpánková et al., 2023; Nagy et al., 2024). In our study, treated animals exhibited significant weight loss, consistent with prior reports linking 4-MU to altered glucose and triglyceride metabolism and enhanced brown adipose tissue thermogenesis (Grandoch et al., 2019; Tsitrina et al., 2023; Nagy et al., 2024). Importantly, rotarod and grip strength performance remained unaffected, and hematological parameters were stable. We noted a mild, non-significant increase in circulating bile acids, aligning with the known pharmacodynamic action of 4-MU on bile acid secretion. Glycosuria occurred without hyperglycemia, suggesting a tubular alteration of glucose reabsorption rather than a systemic effect. Similar findings have been reported elsewhere, with no kidney histopathology, and normalization of biochemical markers post-treatment (Štepánková et al., 2023).

In conclusion, PNN accumulation and neuroinflammation are well-established features of CNS aging that contribute to cognitive decline. 4-MU, a well-characterized inhibitor of HA synthesis with an established pharmacokinetic profile in animals and humans, is clinically approved for biliary cholestasis and under investigation for immune and oncological disorders, making it an attractive candidate for repurposing (Fedorova et al., 2025). Our work characterizes PNN and glial alterations during normal aging and shows that chronic oral 4-MU administration mitigates these changes and increases object recognition memory without evidence of severe adverse effects. These findings support the potential of 4-MU to ameliorate age-related cognitive decline through coordinated functions linked to ECM modulation.

## DATA AVAILABILITY STATEMENT

The datasets presented in this study can be found in online repositories. The names of the repository/repositories and accession number(s) can be found below: https://doi.org/10.5281/zenodo.18223474

## AUTHOR CONTRIBUTIONS

AC: Conceptualization, Data curation, Formal analysis, Investigation, Visualization, Writing – original draft. JD: Conceptualization, Data curation. N.M.V.: Sample extraction. L.M.U.: behavioral analysis. JF: Conceptualization, Funding acquisition, Methodology, Writing – review and editing. PJ: Conceptualization, Funding acquisition, Project administration, Supervision, Writing – review and editing. JK: Conceptualization, Methodology, Supervision, Validation, Writing – review and editing.

